# GARDEN-NET and ChAseR: a suite of tools for the analysis of chromatin networks

**DOI:** 10.1101/717298

**Authors:** Miguel Madrid-Mencía, Emanuele Raineri, Vera Pancaldi

## Abstract

We introduce an R package and a web-based visualization tool for the representation, analysis and integration of epigenomic data in the context of 3D chromatin interaction networks. GARDEN-NET allows for the projection of user-submitted genomic features on pre-loaded chromatin interaction networks exploiting the functionalities of the ChAseR package to explore the features in combination with chromatin network topology. We demonstrate the approach on epigenomic and chromatin structure datasets in haematopoietic cells.

## Introduction

A decade after the first papers about HiC^1^, the techniques to experimentally detect genomic structure at different scales have multiplied. We can now explore 3D chromatin contact dynamics across cell-types^2^, differentiation states^3^, the cell-cycle^4^ and even in single-cells^5^. As the resolution of our datasets improves, the need for new genome visualization options is apparent. We have developed a set of tools for the integration and interpretation of genomics datasets in 3D, based on a network representation of chromatin contacts.

Despite the great advancement in our knowledge regarding chromatin conformation in the last few years, heatmaps are still the most widely used visualization frameworks. These contact matrices are very suited to this data-type, which provides symmetric interaction profiles based on binning of contacts at a specific resolution. This mode of visualization is also ideal to spot two of the main organizational structures in the genome, namely topologically associated domains (TADs)^6,7^, identified precisely as squares on the diagonal of HiC contact heatmaps, and compartments^1^, directly related to the checkerboard patterns that are also easily observable in this representation. Capture HiC is a variation of the HiC protocol that enables studying interactions involving a set of very specific non-abundant regions of the genome, for example gene promoters^8^. Promoter-capture HiC confirmed the existence and importance of long-range chromatin interactions and identified the role of Polycomb in creating a highly interconnected core of developmentally related genes in mouse embryonic stem cells^9^. Long-range interactions can also be detected in contact maps generated by identifying chromatin interactions that are mediated by specific proteins, for example RNA Polymerase II (RNAPII), with the ChIA-PET protocol^10,11^. Hi-C datasets allow the detection of these long-range interactions only when extremely deep sequencing of the libraries is performed, due to the much higher abundance of short-range contacts that saturate sequencing libraries^3^.

One factor which is common to all chromosome conformation capture techniques is the reliance on hybridization and sequencing. Alternative approaches that do not rely on sequencing, and even more so the ones that do not require the bias-prone ligation steps characterising chromosome capture methods, provide us with an independent picture of nuclear organization. For example, different techniques exist for visualizing specific previously tagged chromatin regions by microscopy, leading to the inference of 3D interactions (DNA FISH Oligopaints^12^, Hi-M^13^). Finally, hybrid methods that infer 3D proximity by segregating genomic fragments in different locations in the nucleus allow the inference of 3D contact maps independent of proximity ligation (GAM^14^, SPRITE^15^). There is a clear need to develop frameworks to represent, compare and integrate these different datasets in an intuitive and computationally efficient way. New frameworks to visualize chromatin will help us go beyond the identification of TADs and compartments towards models of gene regulation. We suggest that interpreting chromatin structure as a network is a useful step in this direction.

A few papers have suggested interpreting chromatin structure as a network^16–23^. Despite the processing required to obtain networks from 3C based datasets available at the time, network methods were shown to provide biological insight already in 2012^16^. Thanks to recent improvements in experimental techniques, we are now able to generate meaningful chromatin networks directly from the contact datasets. The network representation is particularly useful when using PCHiC networks, where we can subdivide the whole network into interactions involving only promoters (PP subnetwork) or those involving a promoter and an Other-end (PO subnetwork). This distinction can lead to interesting differences between these two types of genomic contacts, which are characterized by the presence of specific epigenomic features^23^. Representing the genome as a 3D network allows us to find organizing principles that remain hidden when using a linear representation of each chromosome, while also facilitating detection of common structural features such as TADs^24^.

Given the flexibility and efficiency of representing chromatin as a network, we have designed a pair of tools which can be used together or separately for the analysis of chromosome conformation capture data: GARDEN-NET and ChAseR (**Figure 1**). ChAseR is a stand-alone computational tool implemented as an R package, which provides functions to efficiently build and analyse chromatin interaction networks, integrate different epigenomic features on the network, and investigate the relation between chromatin structure and other genomic properties. For example, we have recently introduced the concept of Chromatin Assortativity (ChAs), which allows us to identify whether chromatin with specific features (chromatin marks, binding of transcription factors and replication timing amongst others) tends to form preferential 3D contacts in the nucleus^23,25,26^. ChAseR provides extremely efficient calculations of ChAs and other related measures, including cross-feature assortativity, local assortativity defined in linear or 3D space, and tools to explore these patterns (**Figure 1A**).

**Figure 1:**
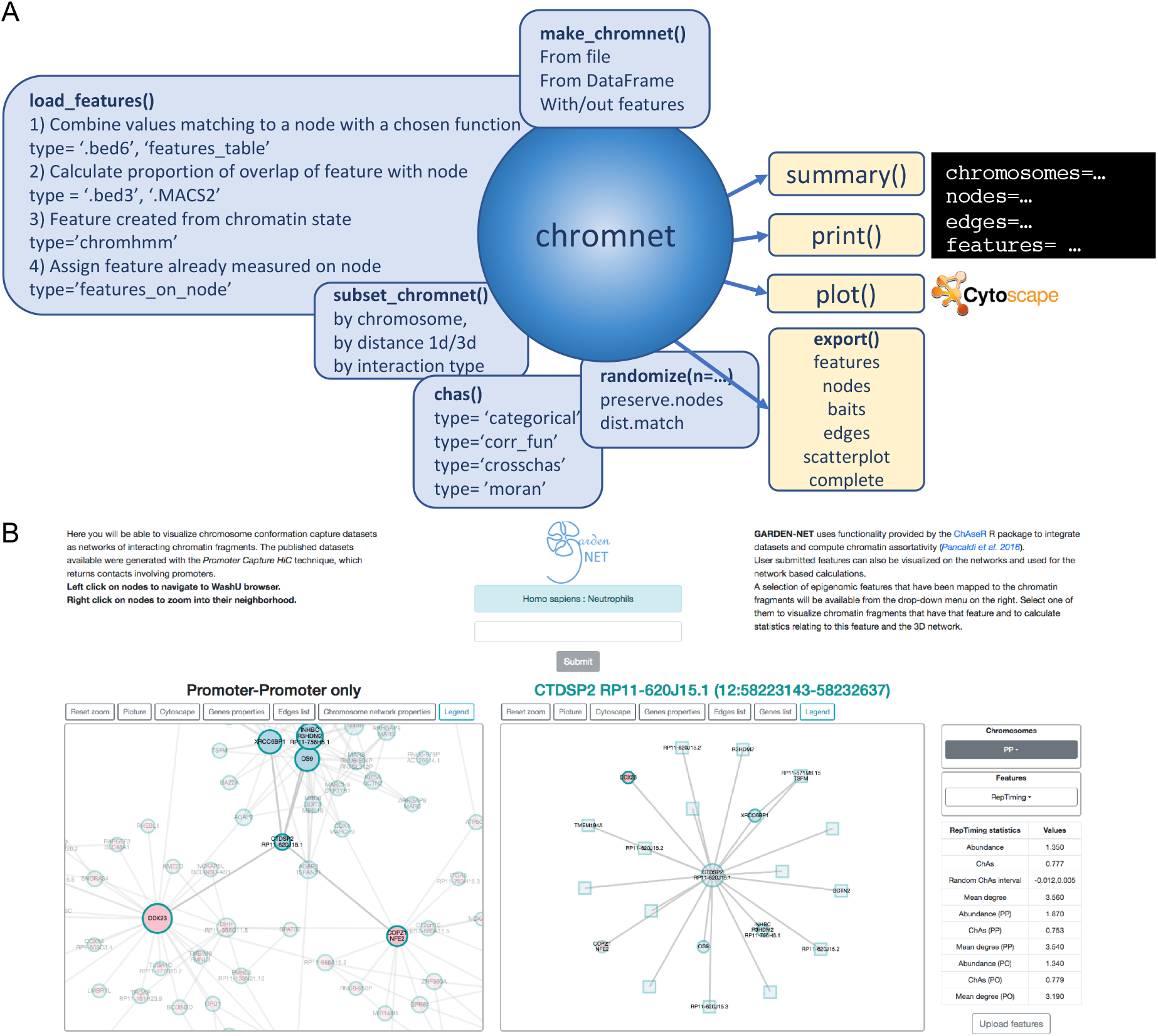
ChAseR and GARDEN-NET. **A**. A schematic overview of the **ChAseR** package showing the main functions that can be applied to the chromnet objects. A file containing a list of edges is made into a chromnet object by make_chromnet. Successively, the user can load a file containing genomic features and use **ChAseR** to assign those features to the relevant nodes in the network. This is done automatically by intersecting the coordinates of the nodes with the coordinates of the features and then calculating a value for each feature on each of the fragments. One can then compute various measures of correlation with **chaser::chas** and check if they are significant with **chaser::randomize**. **B**. A screen-shot from GARDEN-NET showing the different components of the web, including the general chromosome network viewer (left), the neighbourhood network viewer (middle), the menu to choose a chromosome and feature of interest and the table of the network measures related to the selected feature and the button to open the dialogue to upload features (see Supplementary).

GARDEN-NET is a web-based tool that exploits Cytoscape^27^ functionality to allow exploration of pre-loaded chromatin contact networks (>10 PCHiC datasets in human haematopoietic cells^2^ and mESCs^28^) in combination with different pre-loaded epigenomic datasets (histone modifications, expression, DNA methylation^29^ and replication timing) also allowing users to submit features (.bed and other formats accepted). It provides user-friendly visualization and search of networks, calculations of network measures and ChAseR functionalities, including calculation of ChAs.

We demonstrate the use of ChAseR and GARDEN-NET on epigenome datasets from the BLUEPRINT project, in which reference histone modifications, DNA methylation and gene expression were characterised for different haematopoietic cell types in 150 healthy individuals^30^.

We considered contacts identified in more than 10 hematopoietic cell types using PCHiC^2^. Comprehensive epigenomic data was collected for neutrophils, including 4 histone modifications, DNA methylation and expression levels across all individuals, allowing us to test the robustness of our findings in this cell type. For each of the following analyses we performed randomizations, to test the significance of assortativity values (see Supplementary).

The first feature that we examined was tri-methylation of Lysine 27 on histone 3 (H3K27me3), a chromatin mark which is strongly associated to Polycomb repressed chromatin^31^ that was found to be assortative in mESCs^23^. Indeed, H3K27me3 was found to have positive and significant ChAs in neutrophils in both PP and PO subnetworks in most individuals, despite a large variability, confirming the association of Polycomb with 3D chromatin contacts even in differentiated cells (**Figure 2A**).

**Figure 2:**
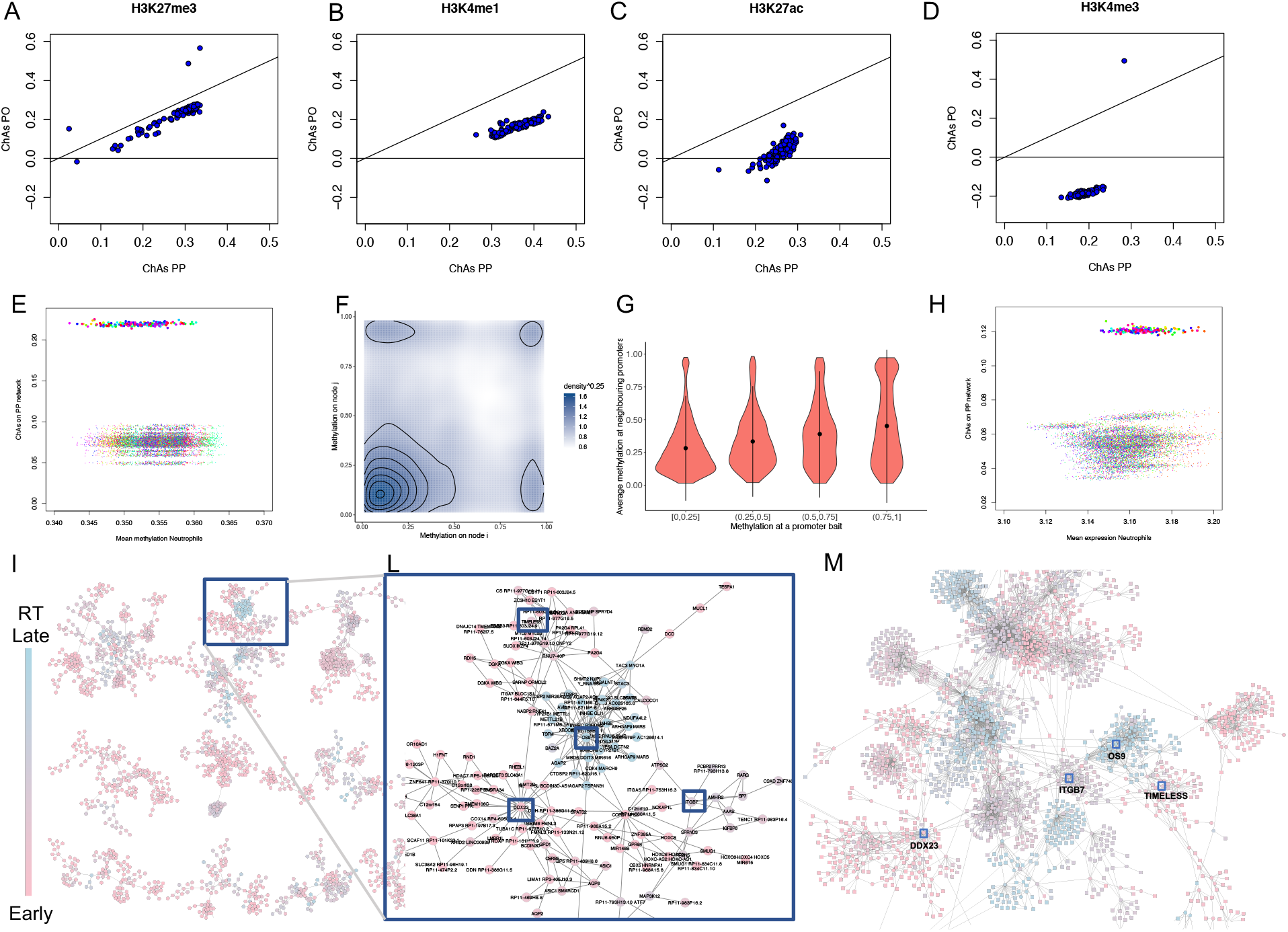
An overview of analyses that can be carried out with ChAseR. **A-D)** Analysis of ChAs of histone modifications on purified neutrophil samples from over 100 individuals^29^ on the PCHiC PP and PO networks for neutrophils^2^ (see also Supplementary Figure S1). **E**) ChAs of DNA methylation from each EPIVAR individual^32^ calculated on the PP PCHiC network^2^ plotted against mean methylation values across all fragments in each individual (filled coloured circles) and 50 ChAs values obtained through distance-preserving randomization in each individual (coloured points). **F**) DNA methylation scatter plots of methylation on nodes connected by edges in the network (methylation averaged over all individuals). **G**) Violin plots showing the distribution of methylation values on neighbouring promoters for promoters with different classes of methylation values. **H**) ChAs of gene expression from each EPIVAR individual plotted against mean expression values across all fragments in each individual (filled coloured circles) and 50 ChAs values obtained through distance-preserving randomization in each individual (coloured points) **I)** Replication timing highlighted by node colour on a subset of the neutrophils PCHiC network (showing only the PP network and where replication timing is taken from values in www.replicationdomain.com for the GM12878 cell line). **L**) Close-up of the region of chromosome 12 inside the blue rectangle in I, representing local accumulation of late replication timing genes (e.g. OS9), surrounded by clusters of early replicating genes (DDX23, TIMELESS) and an intermediate replication timing cluster (ITGB7). **M**) The region of the full PCHiC network corresponding to L, representing bait nodes (circles) and other-ends (squares).

We then examined Lysine 4 mono-methylation on histone 3 (H3K4me1), which has been associated to enhancers and promoters^31^ and was found to be assortative in mESC^23^. H3K4me1 was found to be assortative in neutrophils (**Figure 2B**). ChAs values were higher in the PP than in the PO subnetwork, suggesting that this mark is mostly associated to promoters that are in 3D contact with each other, but it can also be found in some regulatory regions in contact with promoters.

Next, we considered, acetylation on Lysine 27 of histone 3 (H3K27ac), an activation mark which is often encountered on active gene promoters and active enhancers^31^ and was also found to be assortative in mESCs^23^. We see variable levels of H3K27ac assortativity across different individuals, especially in the PO subnetwork with values ranging from −0.1 to 0.1, compared to a range of 0.1 to 0.35 in PP (**Figure 2C**). This suggests that for all individuals, there is a strong tendency for interactions between promoters that have similar levels of activation and in some individuals some regulatory regions in contact with active promoters also have this mark.

Finally, we looked at peaks of trimethylation on Lysine 4 of the same histone 3, a mark associated with active promoters^31^. We observed strong assortativity in the promoter-promoter network and negative assortativity in the PO network (**Figure 2D**), consistent with what was found in mESC^23^ and also with the expected behaviour for a promoter-associated mark. Results for H3K27ac and H3K4me1 could be largely confirmed in monocytes and T cells (**Supp. Figure S2**).

Taken together, these results are consistent with recent reports of the existence of interactions between functional regulatory domains that can be evinced from correlation of histone marks (H3K4me3 and H3k27ac) across a large number of individuals^33^. Moreover, we confirmed an association of chromatin marks to chromatin structure that leads to preferential contacts between chromatin regions with similar chromatin^34^.

Given the strong association between de-methylation at gene promoters and active gene states, we went on to test assortativity of DNA methylation Beta values in the PP subnetwork. Indeed, we found positive values ranging between 0.19 and 0.21. (**Figure 2E**). ChAseR allows for a more detailed investigation of the correlation of features in interacting genomic regions. We can plot the correlation between the value of methylation at one promoter and values of methylation of its neighbouring promoters, separating methylation values in quartiles (**Figure 2F**). Alternatively, we can directly plot the correlation of methylation values in all promoter pairs connected in 3D (**Figure 2G**).

Having observed a strong relationship between chromatin marks related to gene expression and chromatin 3D structure, we predicted that gene expression levels of genes whose promoters establish contacts should be correlated. We confirmed this hypothesis measuring a significant value of assortativity of gene expression on the cell-type specific PP subnetwork in all individuals (**Figure 2H**) and in all three cell types analysed (**Supp. Figure S3**).

Moreover, the general applicability of ChAseR and GARDEN-NET allows us to investigate assortativity of any feature defined along the genome. For example, we found replication timing of genes to be the most assortative genomic feature of the ones analysed, with a value ranging from 0.6 in mESCs to 0.77 in neutrophils (**Figures 1B, 2I-M**).

The assortative patterns of marks related to gene regulation and of the expression levels themselves suggest a relationship between gene regulation and 3D structure at a global level. Moreover, the strong assortativity of replication timing, suggests that replication is also strongly connected to 3D structure, as recently proposed^25^.

## Conclusion

We have presented two complementary approaches to investigate genome-wide datasets in the context of the organization of chromatin inside the nucleus. GARDEN-NET is a chromatin network visualization tool which allows to upload user-defined feature files and project these features on the available 3D chromatin contact networks. The underlying functionalities of GARDEN-NET for the production and investigation of chromatin networks and for the mapping of genome-wide features on them are provided by the ChAseR package, which can also be used on its own. We believe these two tools will empower researchers interested in epigenomics, enabling them to perform both genome-wide and region-specific investigations of their data within the context of 3D genome architecture evinced by chromosome capture experiments, especially when separating different subnetworks is of interest. It should also be noticed that contact networks generated by any other method, including non-sequencing based techniques, can be just as easily used in ChAseR once the experimentally detected contacts are expressed as a list of pairs of nodes corresponding to chromatin fragments.

Despite the power of summarising relationships between 3D structure and specific features with a single correlation coefficient, namely assortativity, the complexity of genome architecture will rarely be fully resolved by this approach. For this reason, we have provided additional tools to more carefully dissect each feature or combination of features in the context of 3D contacts, using extensions of the concept of chromatin assortativity to local and global measures specific for the feature types and suggest scatter plots and boxplots to study the robustness of the correlation at a more granular level.

## Code Availability

GARDEN-NET is available at https://pancaldi.bsc.es/garden-net/ and on GitHub https://github.com/VeraPancaldiLab/GARDEN-NET while ChAseR can be downloaded at https://bitbucket.org/eraineri/chaser/. Code to generate all examples is available at https://github.com/VeraPancaldiLab/ChAseR_demo.

## Acknowledgements

We acknowledge the Barcelona Supercomputing Science for hosting the GARDEN-NET website, Daniel Rico, Biola Javierre, José María Fernández González and Alfonso Valencia for discussions related to the manuscript and Ngoc Tran Bich Cao for testing the R package.

## Author Contributions

Miguel Madrid-Mencía created the GARDEN-NET web-tool, Emanuele Raineri wrote the ChAseR package and documentation. Vera Pancaldi and Emanuele Raineri ran analyses and wrote the manuscript. All authors approved the final version of the manuscript.

## Supplementary Material

### METHODS

#### ChASeR

The ChAseR package provides functions to read in chromatin contact networks and many kinds of data available in bed file format (gene lists, ChipSeq datasets, expression values) and perform the integration of the dataset onto the 3D chromatin structure network. Once the data has been assigned to nodes in the network, ChAseR performs correlation analyses. Specifically, the chas function computes different forms of network correlations: assortativities, cross-assortativities, node by node correlation function, assortativity for categorical features (**see ChAseR vignette attached**).

The value of chromatin assortativity is dependent on network topology, on features abundance (number of network nodes covered by the feature) and also on the relationship between the two (e.g. whether the feature tends to be found in high degree nodes). It is therefore essential to perform randomizations that can suitably estimate the expected value of assortativity for a specific feature to assign a level of significance of the assortativity values identified for a feature of interest. Thus, the package offers various options for performing randomizations.

The simplest option involves label permutation on the network, where labels are given by the values of the feature considered. This type of randomization preserves the number of nodes with a specific feature and also the average value of this feature across the network (in the case of continuous variables). Importantly, when the network contains multiple types of nodes and consequently multiple types of edges, the permutations can optionally preserve the abundance of each feature in each specific node category. For example, when considering PCHiC networks, we subdivide them into P-P and P-O subnetworks and the permutation will preserve abundance in P and O nodes.

Chaser also offers a function for distance matching randomization where the distribution of contact distances in the random network matches approximately that of a real network. In this case the abundance of each feature in each randomization is only approximately preserved. This function is useful for features that have very broad peaks, which are likely to overlap multiple fragments that are linearly close in the genome. In these cases, not preserving distance would produce a distortion of this genomic correlation.

#### GARDEN-NET

Chromatin networks are assembled starting from standard outputs of chromosome capture analysis software. We mostly focused on ‘targeted’ genome-wide techniques such as Capture HiC, ChIA-PET or HiChIP but also standard HiC datasets can be used to create networks after filtering for significant interactions or detection of loops.

To illustrate GARDEN-NET we will here consider data produced by Promoter-Capture HiC which was also the focus of a previous publication^23^. These data formats generally involve the definition of interactions between two genomic fragments and often a score for the strength or confidence in the interaction calling. We thus interpret these interaction files (.ibed) as lists of edges in our network. GARDEN-NET allows to choose specific species and cell types and returns a chromosome-wide visualization of the chromatin contact network on the left panel. The user can then search for a specific region of interest by entering genomic coordinates or the gene name or simply by right-clicking on one of the nodes in the left panel. After the search is activated, the panel on the right will display the neighbourhood of this region/gene while the zoomed in regions are highlighted in the left panel. Alternatively, the user can select to visualize the entire network of only PP interactions and in this case connections between a selected promoter and other ends interacting with it will appear only on the right panel and after the search.

This visualization is based on a web implementation of Cytoscape^27^ and hence visual properties of the graphics can be coupled to the represented dataset. In this case, nodes can be either baits used in the Promoter Capture experiments (circles), or Other Ends (squares). All nodes are annotated with the names of genes that overlap the corresponding genomic fragment (tooltip), and are linked to genome browsers view of the regions (right-click). Bait nodes are additionally labelled with the name of the gene whose promoter was targeted by the capture system. Other Ends often overlap with intronic regions, as shown in the tool tip. The colour of the border of the node denotes the chromosome, to facilitate detection of inter-chromosomal interactions.

A series of tabs above the left-most viewer window allow for resetting of the zoom, downloading of the data and images and loading of a table listing general network characteristics (number of nodes, number of promoter nodes etc.) and network statistics (average degree, size of connected component etc.) for the network represented on the left panel (either a single chromosome or the entire PP network).

The second main component of GARDEN-NET is the mapping of chromatin features on the 3D structure network, which is performed by ChAseR. Depending on the organism and cell type chosen, a list of features is available in the drop-down menu on the right. In the case of mESCs, these are the features that were considered in a previous work^23^ and include histone modifications, binding peaks of different transcription factors and cytosine modifications. For human haematopoietic cells, we currently provide expression, methylation and two histone modifications and expression from the EPIVAR project (average over 150 healthy subjects^32^ and replication timing mapped from the GM12878 cell lines from www.replicationdomain.com. Once one of these features is selected, it will be represented on the network visualization panel with a colour code mapping the node colour to the value. The tool-tip will show the exact value for the feature on the node and the range of feature values in the whole network for comparison.

Upon selection of a feature, a table will also appear below the feature selection drop-down menu with a list of network statistics specific to the selected feature. This includes the proportion of nodes having the feature, their average degree and also the feature chromatin assortativity (ChAs) value, calculated on the whole network (see ChAseR manual about the mapping procedure) with randomized values for reference.

Users are invited to submit their own features through a menu which will require to specify the feature type and provide a feature file (multiple formats are accepted including bed3, bed6, chromhmm) and once the feature is uploaded users will be able to visualize their features on the chromatin network and calculate the related statistics.

### Supplementary Figures

**Supplementary Figure S1:**
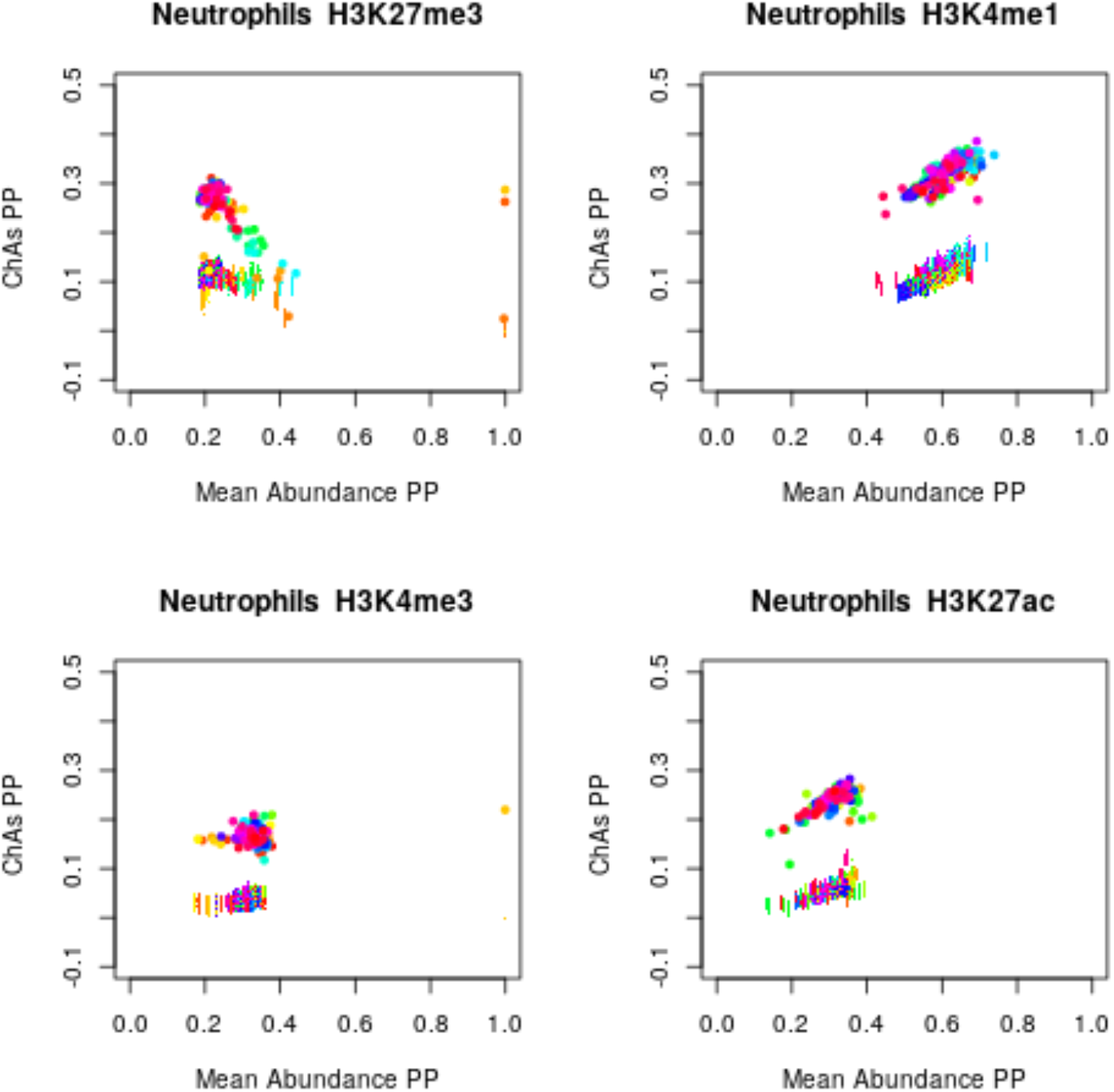
ChAs vs Abundance plots in the PP PCHiC subnetwork for neutrophils for 4 histone modifications (large coloured circles), showing 50 distance preserving randomizations (small coloured dots) (see figure 2A-D).

**Supplementary Figure S2:**
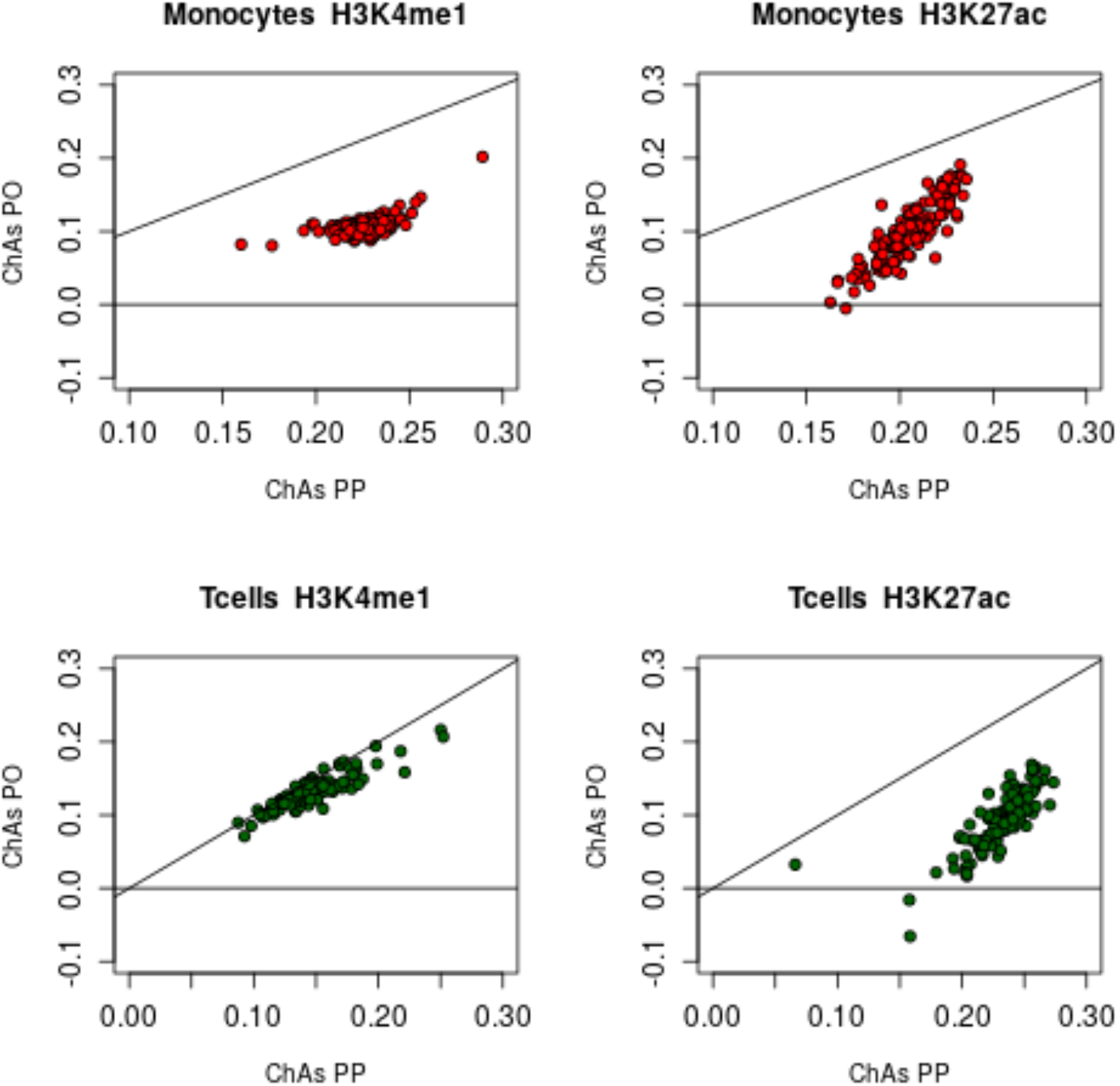
Analysis of ChAs of histone modifications on purified samples from over 100 individuals on the PCHiC PP and PO networks for monocytes (top) and T cells (bottom).

**Supplementary Figure S3:**
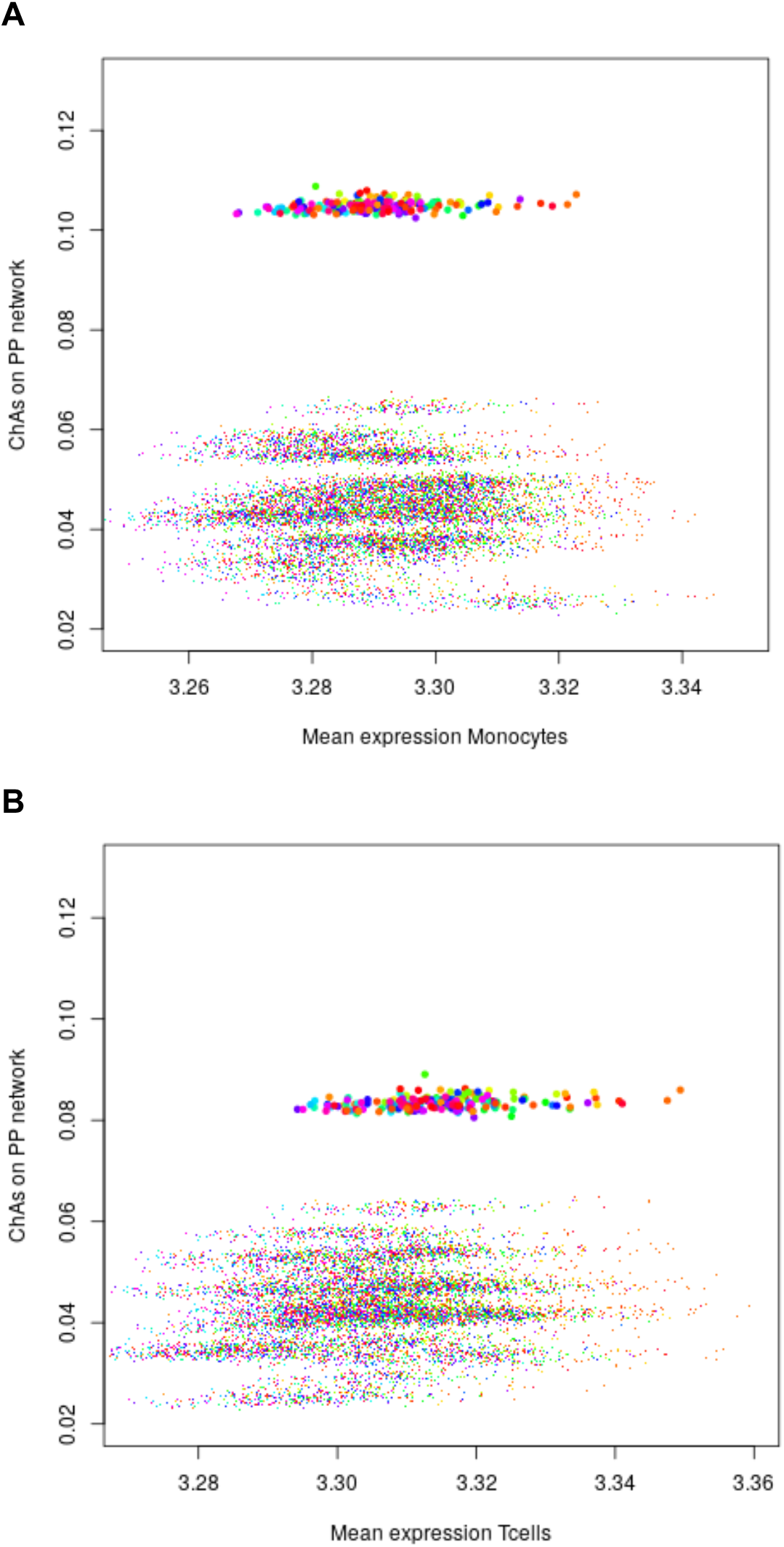
Analysis of ChAs versus mean value of expression. ChAs versus mean expression values in monocytes (A) and T cells (B) showing 50 distance preserving randomizations shown as coloured dots.

## ChAseR: computing correlations in chromatin networks

Emanuele Raineri

2019-07-26

version 0.0.0.9

### Installation

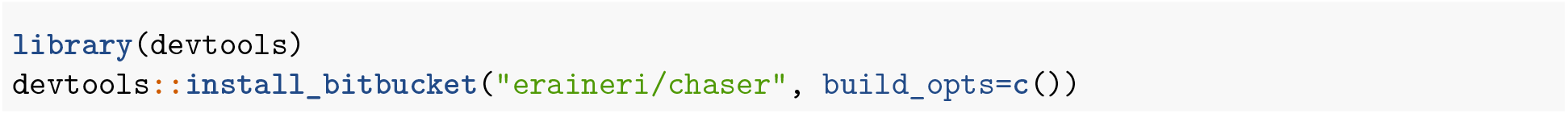

### Introduction

Chromatin networks are a way of representing contacts between regions of the genome which are not necessarily adjacent along the genome but are found to be interating or close to each other in 3D. For example, the interaction between a promoter and an enhancer, or between two promoters of genes that are being co-transcribed. In the past few years they have been analyzed in many different ways (see eg (Pancaldi et al. 2016, Lundberg et al. (2016))). Here we look at the contacts integrating them with other experimental data measured along the genome; for example, chromatin binding peaks measured by ChipSeq. One useful indicator of the relation between the three dimensional structure of the DNA and other features is chromatin assortativity (Pancaldi et al. 2016) i.e. the Pearson correlation of a quantity computed across the edges defined by the 3D contacts.

`ChAseR` is an R package which helps in computing chromatin assortativity efficiently. It tackles four aspects of the computation:

- how to represent the edges of the network starting from an existing dataset (e.g. the result of a promoter-capture experiment (Schoenfelder et al. 2018)
- how to associate to each node in the network one or more genomic features
- how to compute the correlation itself.
- how evaluate the statistical significance of the correlation.

To this purposes chaseR defines a class called `chromnet` and an handful of functions. Objects of class chromnet contain a network and optionally features associated to each node of the network. Note that there is no need for the user to access the chromnet object in any way different from using the functions described below.

There are 6 functions which operate on `chromnet` objects, namely:

- `make_chromnet`
- `load_features`
- `subset_chromnet`
- `chas`
- `export`
- `randomize`

Note that this guide is only a quick introduction to `ChAseR` and does not cover all the possible options of every function. See the online help for more details.

#### creating the network

`make_chromnet` maps a properly formatted data.frame or file to a chromnet object. The data frame must have 6 columns: the first three describe the position of the first node (a genomic region, for example a bait), the second three the position of an other node which interacts with it.

One can also give a file name as argument to `make_chromnet` as in the example below, in which case the data.frame is read from disk. Optionally, it is possible to specify a features matrix.

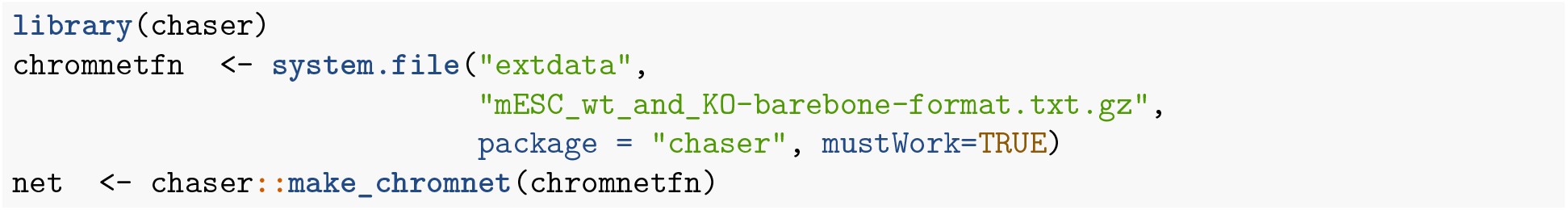

`ChAseR` defines a `summary`, `print` and `plot` functions for the `chromnet` class (the `plot` function requires a running instance of Cytoscape).

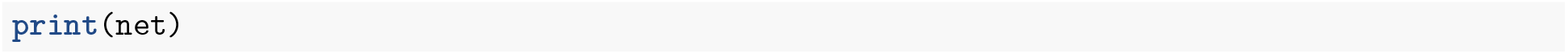

~~~
## nodes: 55855
## edges: 69987
## 22 chromosome(s)
## no features
~~~

The nodes in the network created by chromnet have names which contain the coordinates of the corresponding locus (eg `chr1:10500-11000`).

#### associating features to nodes

Features can be mapped on the nodes using `chaser::load_features`. This accepts the following formats:

- A tsv file which contains the node names in the first column and the features in the remaining columns. This is useful when features are already annotated at the coordinates corresponding to the nodes from, so that there is no neeed for `ChAseR` to perform an overlap. This format correspond to `type=“features_on_nodes”` in the arguments of `chaser::load_features`
- A tsv file (with header) where the first three columns specify a genomic coordinate and the remaining columns are features. One can specify an arbitrary function to modify the features before assigning them on the nodes. (eg one might want to average all the measurements which are assigned to the same node, or take the minimum value, or count how many measurements overlap with the specific node). (`type=“features_table”`).
- bed6 files. Again, one can specify an arbitrary function to process features before assigning them.
- bed3 files. In this case what gets mapped on the node is the fraction of the node occupied by the interval defined by each bed3 line.
- MACS2 output files; again what gets mapped to each node is the fraction of the node occupied by a peak.
- chromhmm files: they produce a feature for each state (what gets mapped to each node is the fraction of the node occupied by the state).

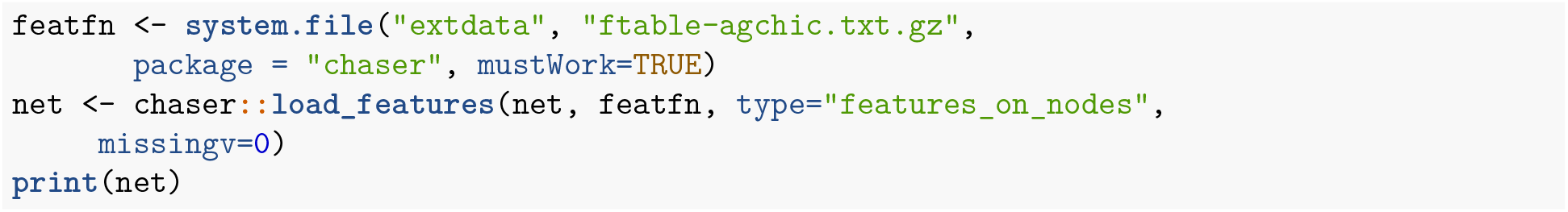

~~~
## nodes: 55855
## edges: 69987
## 22 chromosome(s)
## 78 feature(s)
~~~

#### exporting information from chromnet objects

The function `chaser::export` computes or extracts information from a `chromnet` object. Possible export types are:

- `features` (the default)
- `scatterplot` (to plot a scatterplot of a feature, where each edge is a dot and the values of the features on the two corresponding nodes appear on the x and y axes).
- `nodes` returns a data.frame containing all the nodes in the network
- `igraph` exports the net as an igraph object
- `edges` returns a data.frame containing all the edges in the network.
- `baits` returns the node names of the baits of a promoter-capture experiments, assuming they were stored in the first 3 columns of the data.frame which defined the network.

#### extracting subnetworks

Through `subset_chromnet` one can create subnetworks for example by chromosome

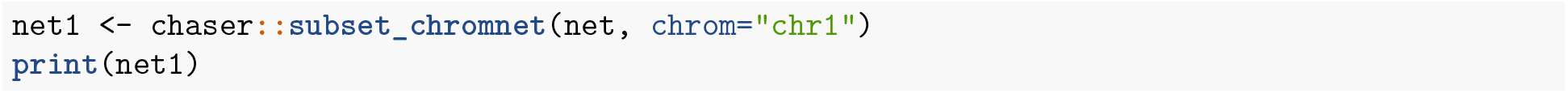

~~~
## nodes: 3120
## edges: 3362
## 1 chromosome(s)
## 78 feature(s)
~~~

or one can select a special set of nodes and only consider edges joining nodes which belong to the set as in the following snippet:

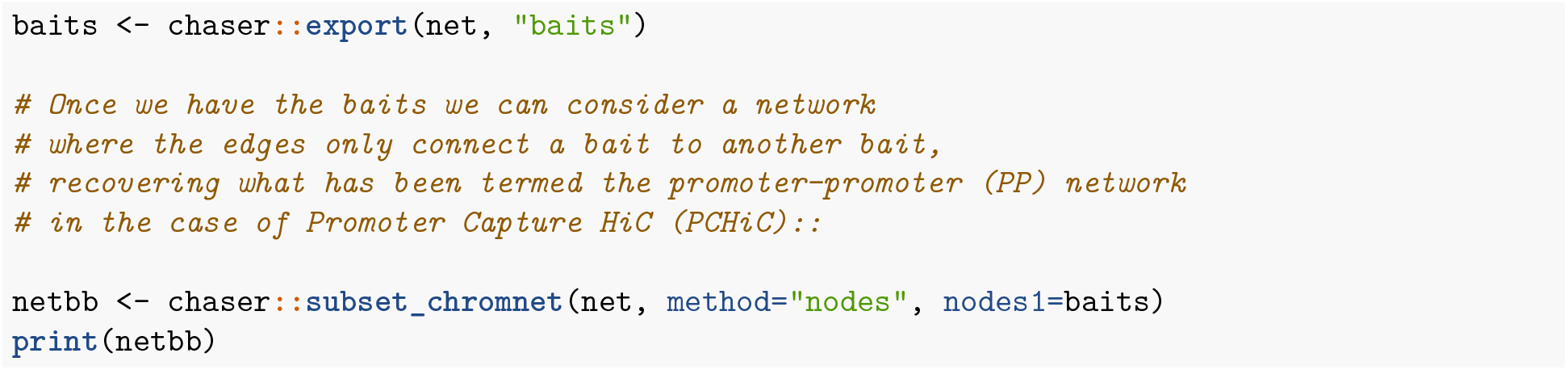

~~~
## nodes: 13099
## edges: 18795
## 21 chromosome(s)
## 78 feature(s)
~~~

Similarly one can construct a network where edges exist only between a bait and an *other end*, which corresponds to the PO network in the case of PCHiC.

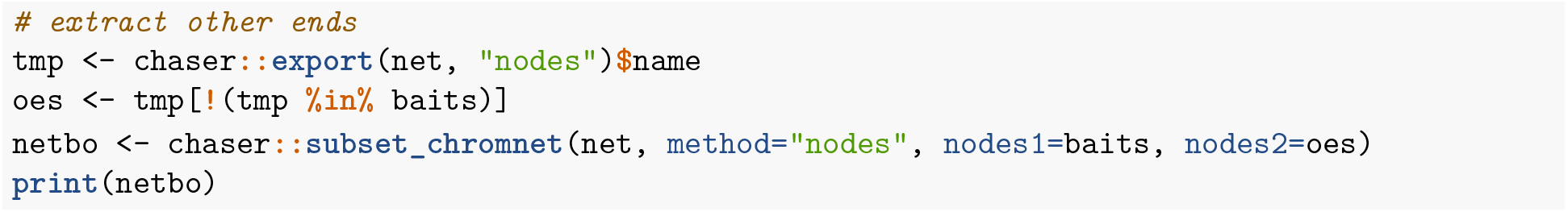

~~~
## nodes: 54123
## edges: 51192
## 22 chromosome(s)
## 78 feature(s)
~~~

It is also possible to select a node and consider the subnetwork formed by all the nodes close (along the sequence) to it.

Here we compute the assortativity on chr1 on windows of 10Mb along the genome. The plot on the bottom is the number of nodes present in each window.

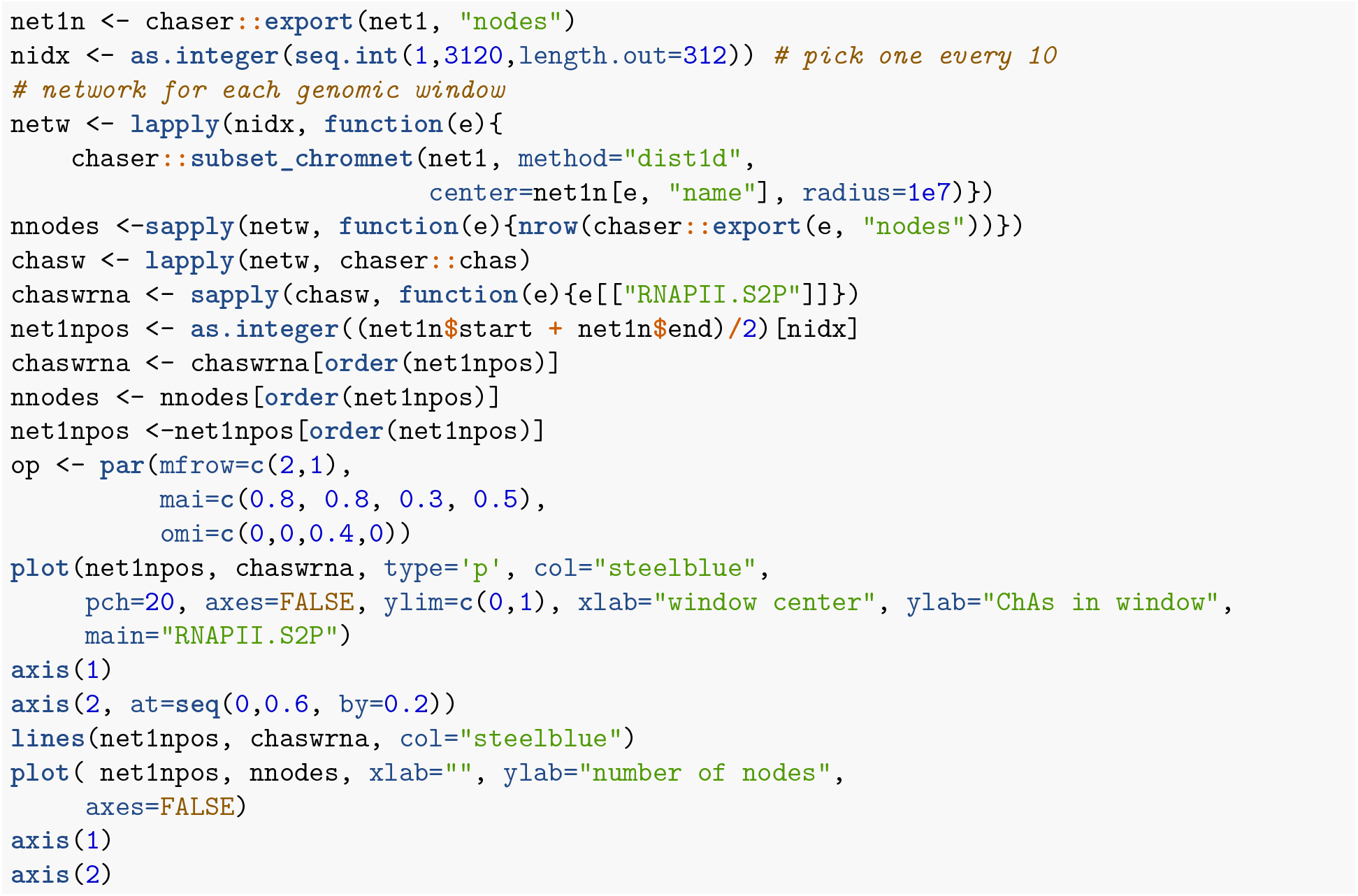
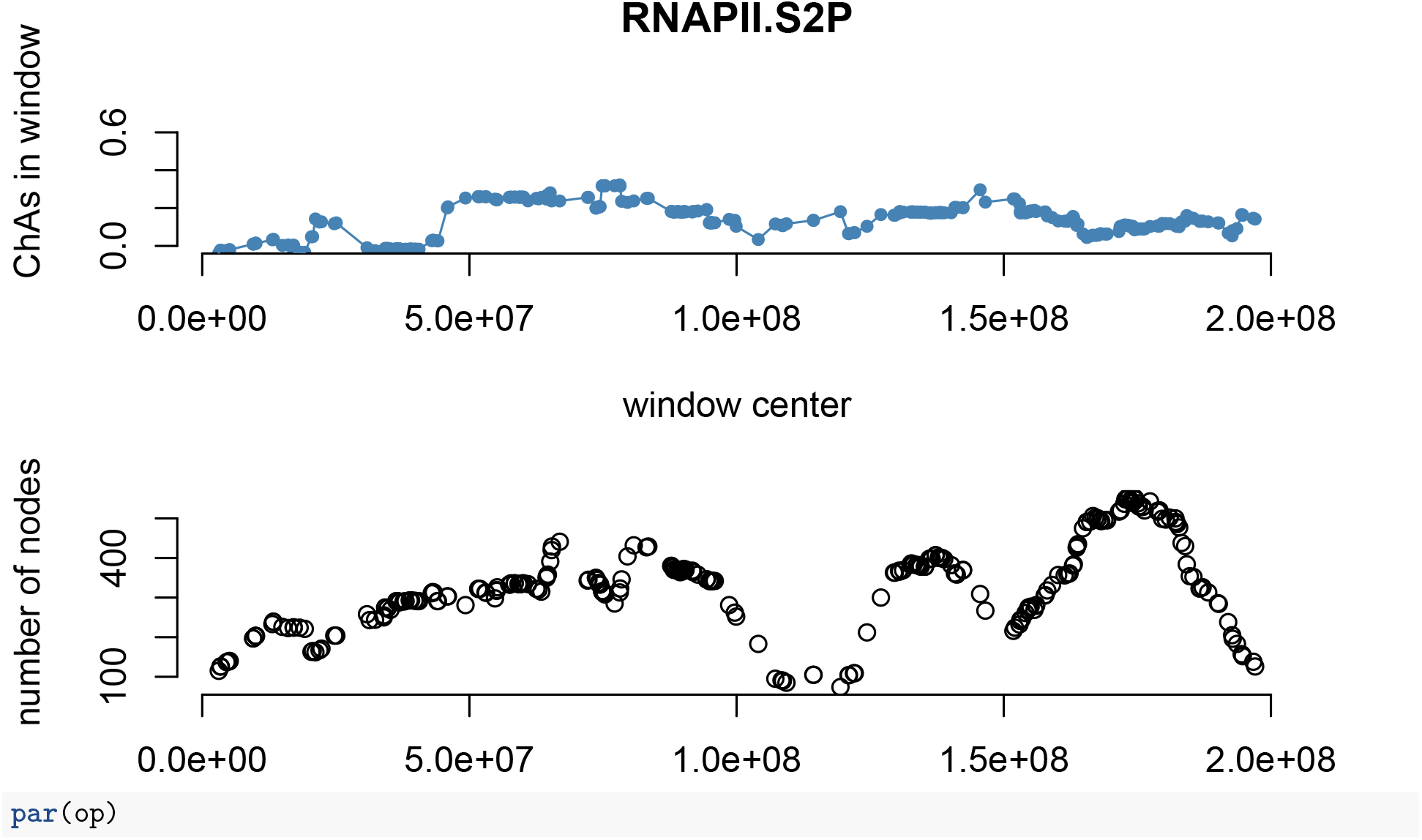

another option of `subset_chromnet` allows one to select a subnetwork based on three dimensional distance (measured in number of edge hops from one node to another on the contact network).

#### computing correlations

The `chaser::chas` function computes different forms of network correlations, depending on the method specified by the user:

- a vector of assortativities, one for each feature mapped onto the network
- a matrix *k* × *k* containing cross-assortativities between all possible pairs of features. This includes the assortativities of each feature with itself, which lie on the diagonal of the matrix.
- a vector containing the Moran index for each feature
- a matrix *n* × *k* which associates to each node the mean value of each feature computed on all its neighbours.

One useful plot is assortativity versus abundance:

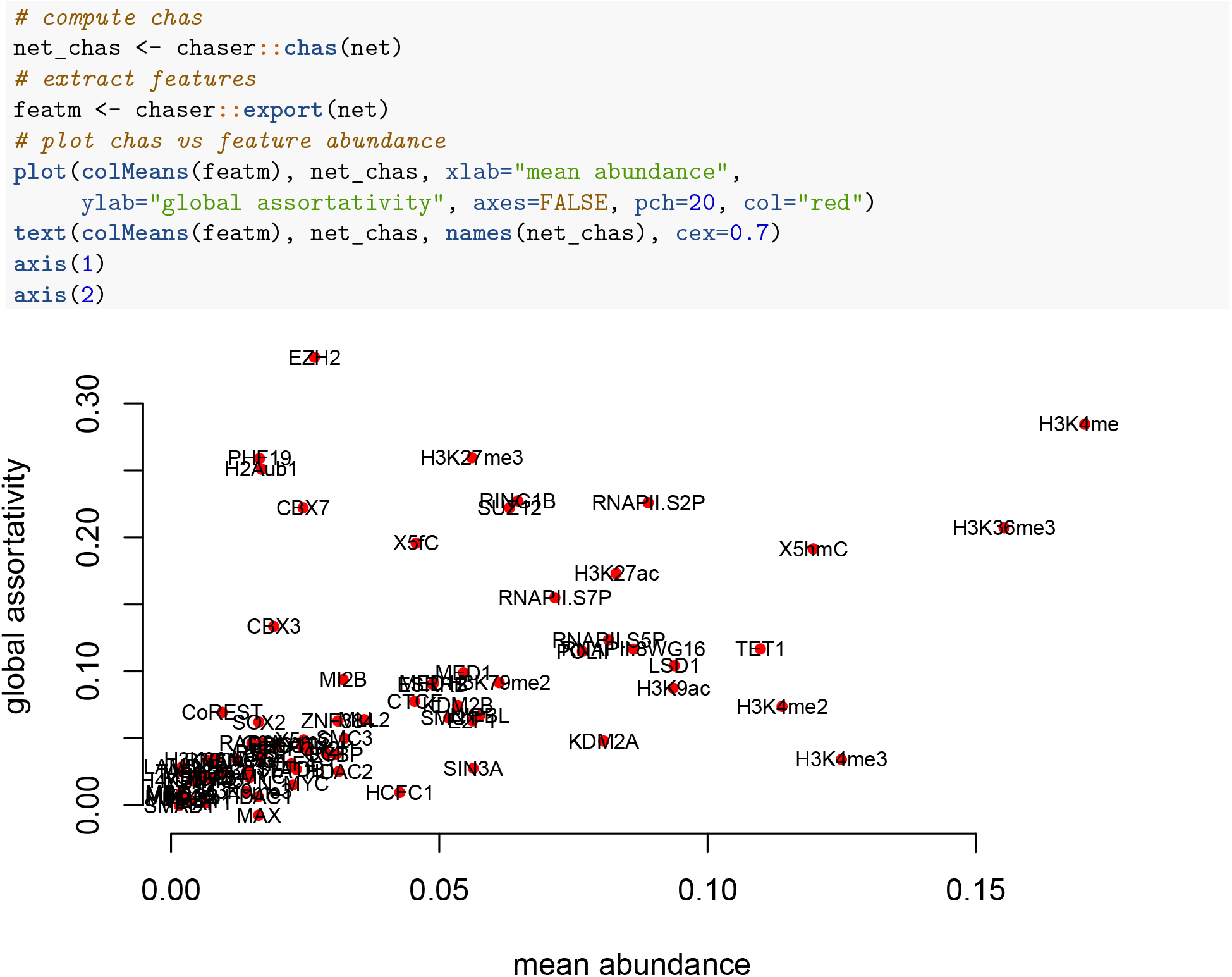

Here we give an example of how to use the `corr_fun` option of `chas` to look at correlations in a more granular way than by giving a single number for the whole network: we plot for each node the abundance at that node and the average abundance of the neighbours.

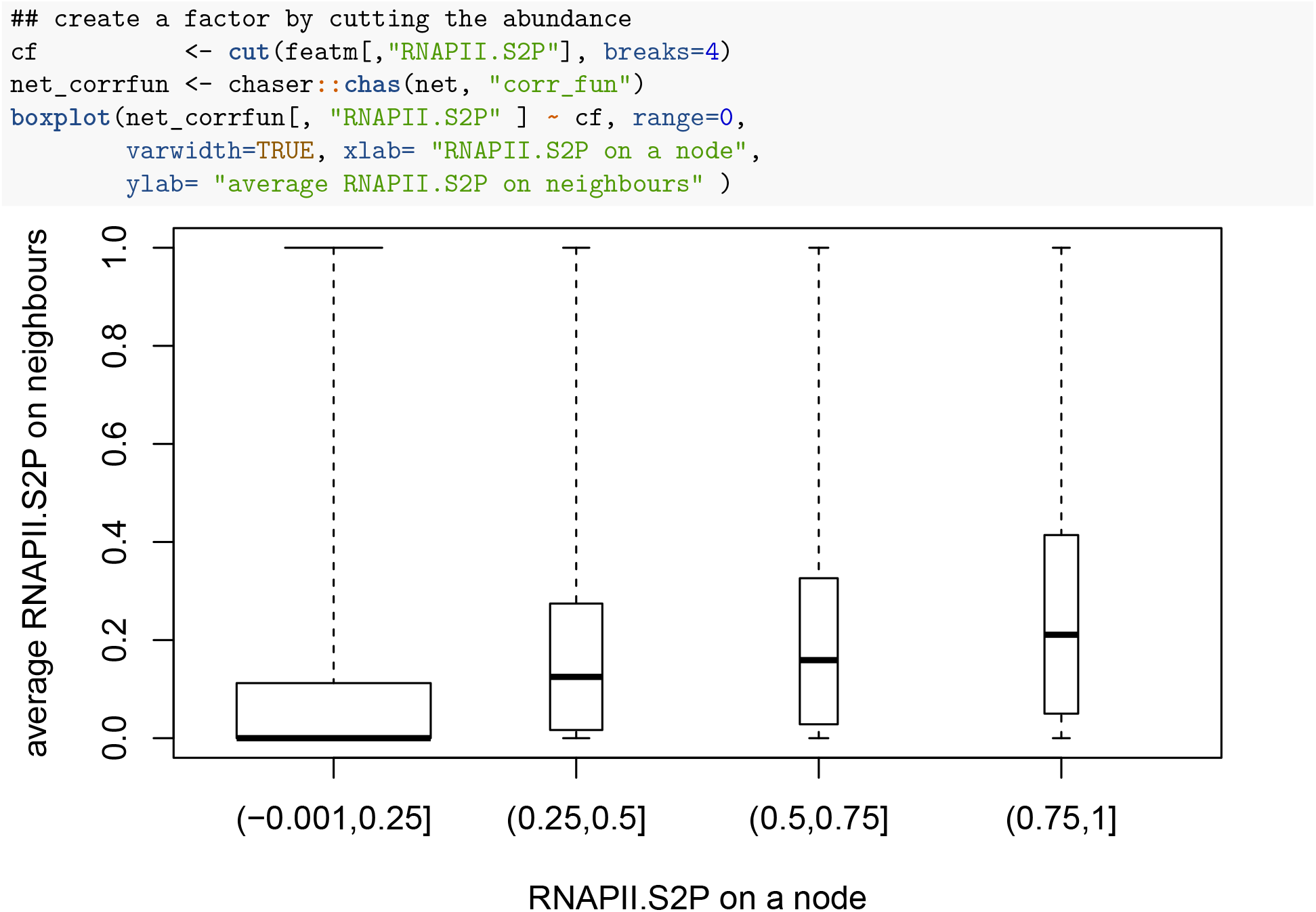

#### A note on assortativity

In what follows, let’s assume that the matrix *F* containing the features has n rows (corresponding to *n* nodes) and *k* columns. Let *A* be a symmetric *n* × *n* matrix, the adjacency matrix of the network. We can safely assume *A* (although not *F*) to be sparse.

As said before, chromatin assortativity is the correlation computed across the edges; i.e. the random variables being paired are the nodes at the end of the same edge. Let *x_i_*(*x_j_*) be the value of a feature at node *i*(*J*). The Pearson correlation is given by

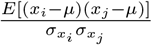

where *μ* is the mean of the feature across all nodes, weighted by the number of edges they take part in 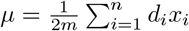 where *d_i_* is the degree of node *i* and *m* is the number of edges. similarly 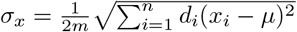 if we normalize each column of *F* by subtracting its mean and dividing by its *σ* we obtain a matrix 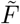 such that the assortativity (as a matrix *k* × *k*) is given by:

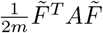

#### randomize

##### vanilla randomization

one can randomize the network by redistributing the features in the nodes at random, before computing the correlation, as in the following example

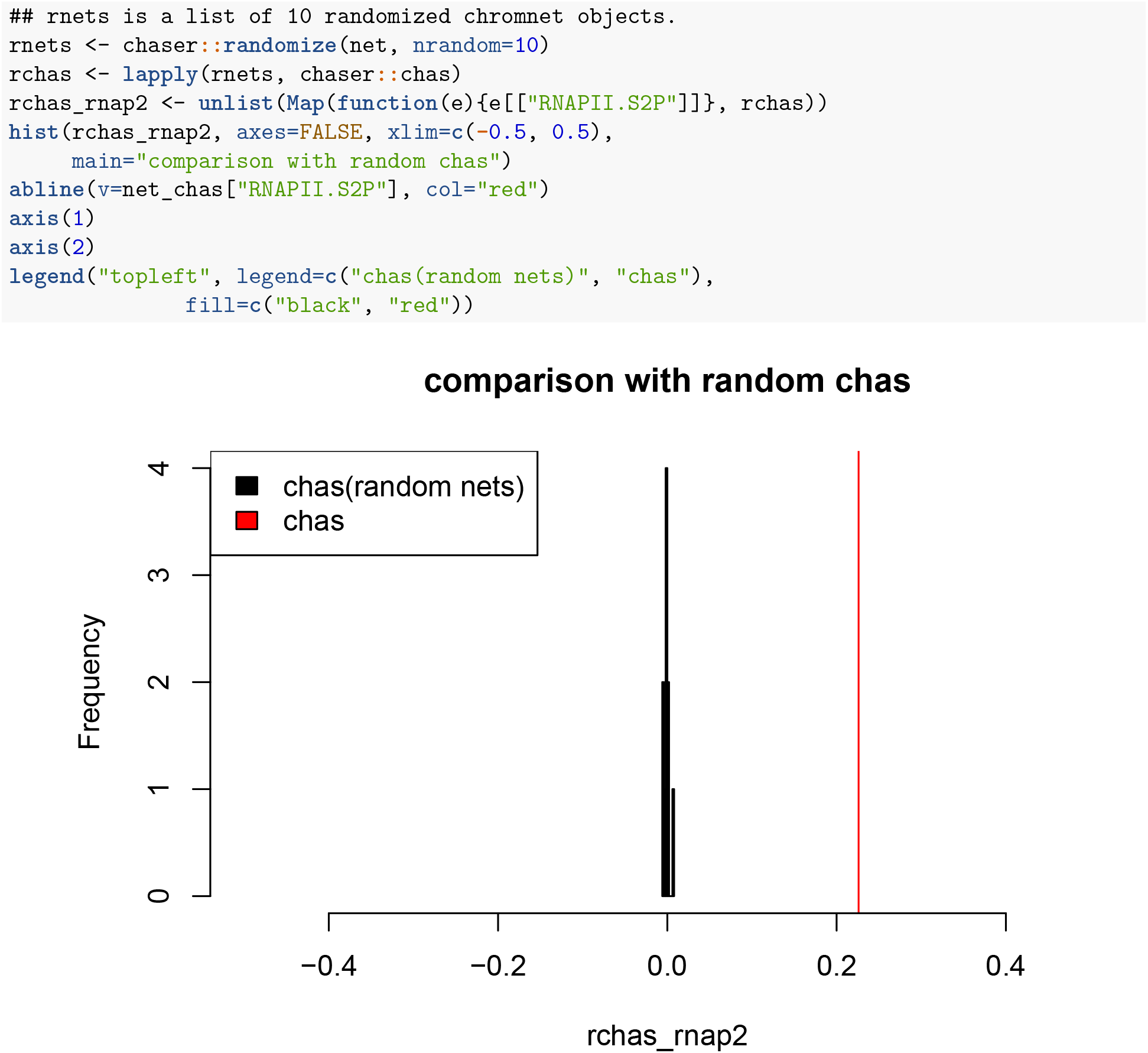

##### randomization with a set of invariant nodes

The randomizaton can be more subtle, for example one can divide nodes into two non overlapping groups and features will be redistributed only inside each group.

In the following examples we use the group baits (and implicitly the group *other ends*) to this purpose.

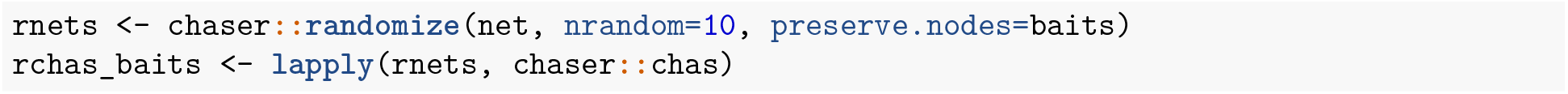

##### randomization preserving distances

Finally (by setting `dist.match=TRUE`) the randomization preserves (approximately) the distribution of the genomic distances spanned by the connected nodes.

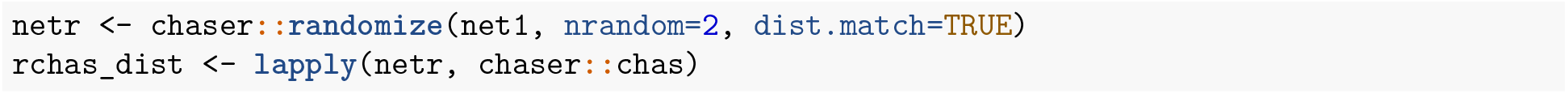

The code above creates a list `netr` of length 2 (in order not to slow down the compilation of this vignette); each element of the list `rchas_dist` is an array of 78 assortativities computed on randomized networks.

We can compare the distribution of distances in the true network with the distribution of distances in the randomized network:

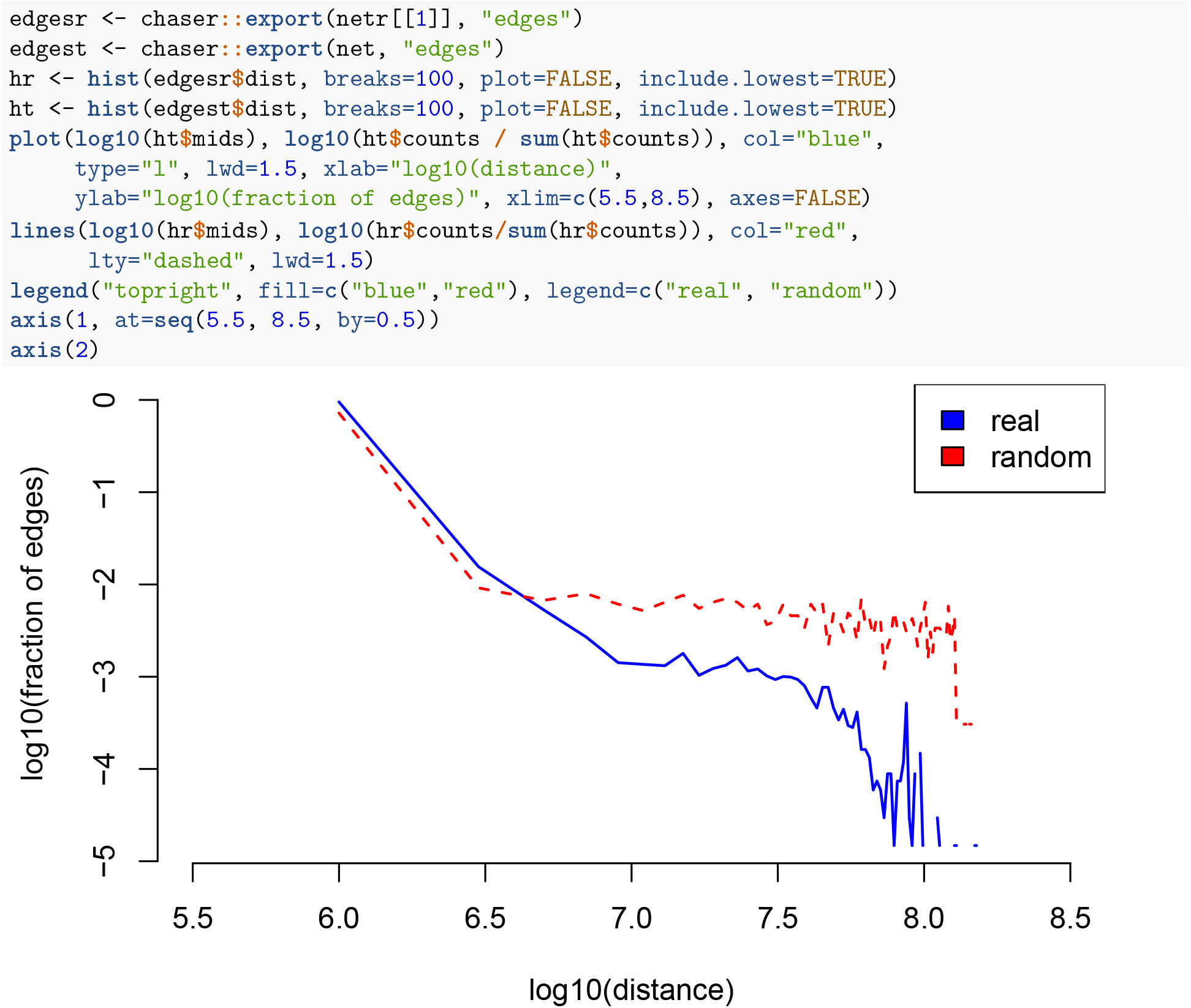

The algorithm which computes random networks with the same distance distributions as the original network works as follows. I assume here that I want to get a distance distribution with approximately the same quartiles as the original network, but the description can be easily extended to a different set of percentiles. Also, let the intervals *I*_1_,*I*_2_,*I*_3_,*I*_4_ be the fourth quartiles of the true distance distribution and *q*_1_, *q*_2_, *q*_3_, *q*_4_, *q*_5_ the corresponding endpoints.

Note that the algorithm randomizes chromosomes one at the time and that the randomized network will not contain interchromosomal contacts.

1. set `i <− 1` (this counts the number of extracted edges)
2. select a node at random 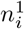
3. select which interval of distances to extract from as `q <− i mod 4 + 1`
4. find all the nodes whose distance from *n_i_* lies in *I_q_*
5. if step 4 return no nodes go to step 2
6. select at random one of the nodes found at step 4 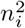
7. store the edge 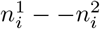
8. set `i <− i + 1`
9. if `i` is bigger than the number of edges in the original network, exit
10. go to step 2.

The actual code includes some heuristic evaluations to avoid attempting randomization when it is not feasible (for example because there are very few edges).

## Discussion

In the diagram below we illustrate a typical `ChAseR` workflow. A file containing a list of edges is made into a chromnet object by `make_chromnet`. Succesively, the user can load a file containing genomic features (in the figure we hint at H3K27ac peaks) and use `ChAseR` to assign those features to the relevant nodes in the network. This is done automatically by intersecting the coordinates of the nodes with the coordinates of the features. One can then compute various measures of correlation with chas and check if they are significant with `randomize`.

---

**Figure 1:**
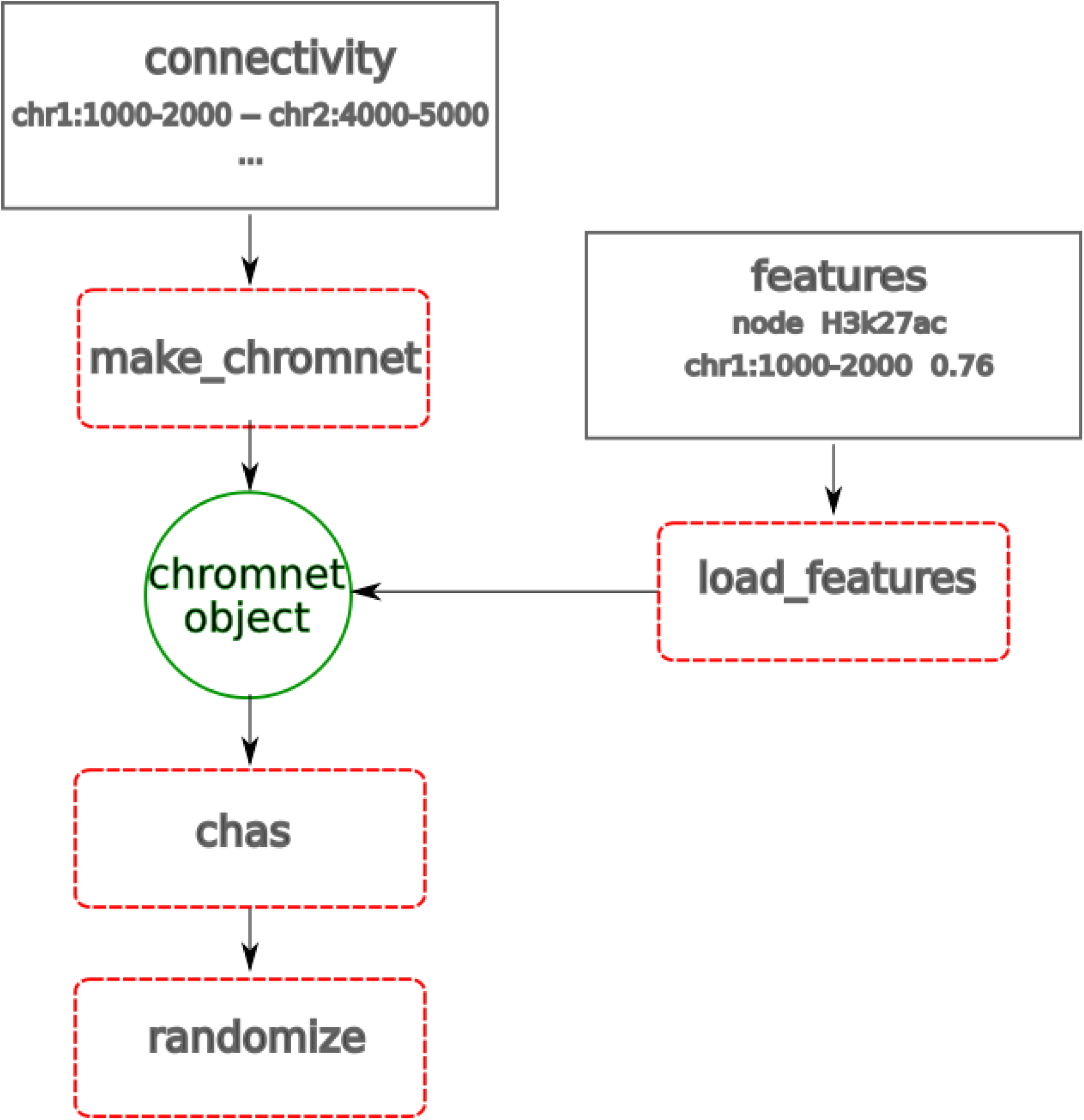
A ChAseR workflow

